# Comparative genomic analysis and characterization of *Staphylococcus* sp. AOAB, isolated from a notoriously invasive *Mnemiopsis leidyi* gut revealed multiple antibiotic resistance determinants

**DOI:** 10.1101/2021.01.11.426108

**Authors:** Richard M. Mariita, Mohammad J. Hossain, Anthony G. Moss

**Affiliations:** College of Science and Mathematics, Biological Sciences Department, Auburn University, AL 36849, USA; Microbial BioSolutions, 33 Greene Street, Troy, NY, 12180 USA; The Department of Biological Chemistry, The Johns Hopkins University School of Medicine, 725 N. Wolfe Street, Baltimore, MD 21205-2185, USA

## Abstract

Here, we describe the isolation and characterization of a coagulase-negative, vancomycin and oxacillin-susceptible novel bacterium of the genus *Staphylococcus. Staphylococcus* sp. strain AOAB was isolated from the stomodeum (gut) of the *Mnemiopsis leidyi* from Mobile Bay, Alabama USA. A polyphasic taxonomic approach comprised of phenotypic, chemotaxonomic and genotypic characteristics was used for analysis. The dominant respiratory quinone detected was MK-7 (100%). Major cellular fatty acids were anteiso-C_15:0_ (40.52%), anteiso-C_17:0_ (13.04 %), C-_18:0_ (11.53%) and C-_20:0_ (10.45%). The polar lipid profile consisted of glycolipid, phospholipid, phosphatidylglycerol and diphosphatidylglycerol. Although strain AOAB had a 16SrRNA gene sequence similarity of 99% with *S. warneri* SG1, *S. pasteuri, S. devriesei* KS-SP_60, *S. lugdunensis* HKU09-01, *S. epidermidis* RP62A, *S. haemolyticus* JCSC1435 and *S. hominis* DM 122, it was be distinguished from those species based on Multi-Locus Sequence Analysis (MLSA) using 6 marker genes (16S rRNA, *hsp60, rpoB, dnaJ, sodA* and *tuf*). MLSA revealed strain AOAB to be closely related to *S. warneri* SG1 and *S. pasteuri* SP1 but distinct from two hitherto known species. These results were confirmed by Average Nucleotide Identity (closest ANI of 84.93% and 84.58% identity against *S. warneri* SG1 and *S. pasteuri* SP1 respectively). *In-silico* DNA-DNA hybridization was <70% (33.1 % and 32.8% against *S. warneri* SG1 and *S. pasteuri* SP1 respectively), which further confirmed that the strain was a potential novel *Staphylococcus* species.

## Introduction

*Mnemiopsis leidyi* (Phylum Ctenophora) is a predatory gelatinous zooplankter endemic to the Western Atlantic, notorious for its invasion of all of the European enclosed and coastal seas, including the North Sea, the Baltic, the Black and Caspian Seas and the southern Adriatic and Mediterranean seas. *M. leidyi* is similarly notorious for its ability to alter the composition of native plankton communities (Delpy, Pagano, Blanchot, Carlotti, & Thibault-Botha, 2012; Jaspers et al., 2012; Lucic et al., 2012). *Mnemiopsis* blooms have been found to coincide with profound environmental perturbation (Purcell, 2012). Much like other zoonotic organisms in exotic habitats (DeLong, 2014), they also have been revealed to be vectors for microbial assemblages (Daniels & Breitbart, 2012; Hao, Gerdts, Peplies, & Wichels, 2015; Moss, Estes, Muellner, & Morgan, 2001). Although ctenophores are known to be parasitized by amoebae, dinoflagellates, sea anemones and bacteria (Daniels & Breitbart, 2012; Hammann, Moss, & Zimmer, 2015; Hao, Gerdts, Peplies, & Wichels, 2015; Moss, Estes, Muellner, & Morgan, 2001), there is no study that has isolated individual ctenophore gut bacteria for characterization.

This study used phenotypic, genotypic and phylogenetic analysis to characterize a *Staphylococcus* isolate AOAB. Previous studies have isolated novel staphylococci species from marine waters (Gunn & Colwell, 1983), fresh water (Hess & Gallert, 2015), marine crustaceans (Faghri, Pennington, Cronholm, & Atlas, 1984), domesticated animals (Yamashita et al., 2005) and human hosts (Trulzsch et al., 2007), among other sources.

## Materials and Methods

Ctenophore gut (stomodeum) samples were collected from animals from Dauphin Island Marina, Mobile Bay, Alabama. Sterile toothpicks were inserted into the gut from the oral end and gently rotated, in order to collect a mucus-rich sample without damage to the stomodeal lining. Tips containing the mucoid gut samples were transferred into 1.5 mL tubes, stored on ice and transported to the laboratory for enrichment.

Enrichments were done using a modified protocol from Anacker and Ordal. Modifications were guided by Figueiredo et al. and Pilarski et al. (Anacker & Ordal, 1955; Figueiredo et al., 2005; Pilarski, Rossini, & Ceccarelli, 2008) (Supplementary Material 1). After an initial stage of enrichment, cultures isolated and maintained using mannitol salt agar (Thavasi, Aparnadevi, Jayalakshmi, & Balasubramanian, 2007) and LB agar (Sigma Aldrich).

Phenotypic tests were carried out following minimum recommended standards to aid in discriminating species (Freney et al., 1999) using using carbohydrate fermentation with the API CH50 systems (bioMe’rieux). Optimal growth was obtained at 37°C after 24-48 hrs.

Chemotaxonomic analyses were also carried out to characterize phenotypes. To assign genus and confirm coagulase test of the isolate, occurrence of fatty acids ai-C15:0, i-C15:0, i-C17:0, ai-C17:0 and menaquinone (MK) in the cytoplasmic membrane was investigated as recommended (Freney et al., 1999; Heß & Gallert, 2015; B. J. Tindall, Rosselló-Móra, Busse, Ludwig, & Kämpfer, 2010). The Analysis for respiratory quinones was carried out by first separating them from other classes using thin layer chromatography on silica gel (Macherey-Nagel Art. No. 805 023), using hexane:tert-butylmethylether (9:1 v/v) as solvent. Menaquinones were then removed from the plate and analyzed using HPLC fitted with a reverse phase column (Macherey-Nagel, 2 mm x 125 mm, 3 µm, RP18) with methanol: heptane 9:1 (v/v) being used as the eluant. Polar lipids were extracted from 100 mg of freeze dried bacterial cells using chloroform: methanol: 0.3% aqueous NaCl mixture 1:2:0.8 (v/v/v), separated using two-dimensional silica gel TLC (Macherey-Nagel Art. No. 818 135) and detected using methods described by Tindall et al. (Brian J. Tindall, Sikorski, Smibert, & Krieg, 2007) (DSMZ Identification Services, Braunschweig, Germany). Cellular fatty acid composition for strain AOAB was determined using gas chromatography (chromatograph was fitted with a 5% phenyl-methyl silicone capillary column) using Sherlock MIS (MIDI Inc, Newark, USA) system. Using MIDI software package (MIDI Inc, Newark, USA), the fatty acid composition data was also used for clustering using Euclidean method for differentiation of isolate from closest species.

Antimicrobial susceptibility was performed using a standard disk diffusion method, following cutoff ranges as outlined by the National Committee for Clinical Laboratory Standards (NCCLS) (Miller et al., 2003). Isolates were inoculated using spread plating with 100 µL of inoculum at log phase from LB broth onto LB agar before impregnating with antibiotic discs. Zones of inhibition (ZOI) were used to determine the ability of the antibiotics to inhibit the growth. The antibiotics used are shown in Table 1. Results were interpreted using commonly accepted zone breakpoints for *Staphylococcus* (Howe & Andrews, 2012). Control discs were sterile 6mm discs without antibiotics.

**Table 1:** Antibiotic susceptibility: ZOI (Zone of inhibition) as measured (in mm) from the edge of disc to edge of inhibition zone (Kaushik, Kessel, Ratnayeke, & Gordon, 2015). Resistant (R) and susceptible (S) assigned based on previously defined breakpoints (Howe & Andrews, 2012). Below the breakpoints, a given isolate is resistant. Antibiotics with unrevised break points are indicated as dash (-). Experiments done in triplicate.

| Antibiotic | ZOI 1 | ZOI 2 | ZOI 3 | Avg. | R/S | Breakpoint (mm) |
| --- | --- | --- | --- | --- | --- | --- |
| Bacitracin 0.04 iu (A) | 6 | 6 | 6 | 6 | R | >14 |
| Nalidixic acid 30 µg (NA30) | 8 | 6 | 7 | 7 | R | >19 |
| Ciprofloxacin 5 µg (Cip5) | 13 | 15 | 14 | 14 | R | >21 |
| Kanamycin 30 µg (K30) | 15 | 14 | 14 | 14.3 | R | >18 |
| Penicillin (P10 iu) | 15 | 15 | 15 | 15 | R | >25 |
| Ampicillin 10 µg (AM10) | 15 | 17 | 17 | 16.3 | R | >26 |
| Vancomycin (VA 30 ug) | 19 | 18 | 18 | 18.3 | S | >12 |
| Tetracycline 30 µg (TE30) | 24 | 23 | 23 | 23.4 | S | >20 |
| Amoxicillin/Clavulanic acid 20/10 µg (AMC30) | 24 | 23 | 28 | 25 | S | >25 |
| Chloramphenicol 30 µg (C30) | 30 | 27 | 28 | 28.3 | S | >15 |
| Novobiocin 30 µg (NB30) | 35 | 35 | 35 | 35 | S | >16 |
| Oxacillin 1µg (OX1) | 20 | 19 | 18 | 19 | S | >15 |
| Control: Polymyxin B ( PB300 iu) | 6 | 6 | 6 | 6 | R | NA |

R=resistant; S=susceptible

Genomic DNA was isolated using the CTAB protocol (Andreou, 2013) with minor modifications that included the use of 0.5 mm silica beads (Biospec Products, Inc. Cat. No. 110791052z), shaken using a specialized MO BIO Vortex-Genie^R^ 2 (MO BIO Laboratories). DNA purity was checked using a NanoDrop reader (ND-2000, NanoDrop Technologies, Wilmington, DE, USA) and precision-quantified using Qubit HS reagents (Life Technologies). The DNA template concentration was adjusted to 5ng prior for use in touchdown PCR reactions. Amplification for most of the 16S rRNA gene was achieved using the universal primers 63f (5′-CAG GCC TAA CAC ATG CAA GTC-3′) and 1387r (5′-GGG CGG WGT GTA CAA GGC-3′) as described by Suriyachadkun *et al*. (Suriyachadkun et al., 2009).

Contents of a 25 µL PCR mixture included 12.5 µL of EconoTaq Plus Green 2X Master Mix (Lucigen), 0.5 µL of 20 µM of forward and reverse primers, 10.5 µL water and 1 µL of 5 ng µL^-1^ genomic template DNA. PCR was carried out using ‘touchdown’ conditions: initial denaturation was at 95 °C for 5 min, followed by 20 cycles at 95 °C for 1min 61°C for 45 sec, and extension at 72 °C for 90 sec. The touchdown method was followed by another 30 cycles at 95 °C (1min) 51 °C (45 sec) 72 °C (90 sec). The final extension was done for 7 min at 72 °C.

The study also targeted a 370 bp *tuf* gene which is well established for *Staphylococcus* taxonomy (Martineau et al., 2001). Primer selection and melting temperature determination was done using an *silico* PCR simulator (http://insilico.ehu.es/PCR); (San Millan, Martinez-Ballesteros, Rementeria, Garaizar, & Bikandi, 2013). The PCR mixture included 0.4 µM of each *Staphylococcus*-specific primp0ers (*tuf*-F (TStaG422) 5′-GGC CGT GTT GAA CGT GGT CAA ATC A-3 and *tuf*-R (TStag76) 5′TIA CCA TTT CAG TAC CTT CTG GTA A-3′) (Tm 59.3°C). The PCR reagent ratios were similar to those used in the 16S rRNA gene amplification. Touchdown PCR conditions were as previously described (Martineau et al., 2001) with modifications that included 5 min at 95°C, 30 cycles of 30 s at 95°C, 30 s at 55°C and 45 s at 72°C, and final extension for 7 min at 72°C. Amplified PCR products were purified using QIAquick (Qiagen, Maryland, USA) purification kit following manufacturer’s instructions prior to sequencing using ABI 3100 DNA Genetic Analyzer at Auburn University’s Genomics & Sequencing Laboratory (GSL).

Genomic DNA of the isolate was used for library construction. We used the Agilent 2100 BioAnalyzer (Agilent Technologies, USA) to perform size fractionation and quantification of DNA. DNA was then fragmented and libraries prepared using Nextera XT (Illumina) according to manufacturer’s protocol before being run on an Illumina Miseq (for 2 x250 bp paired-end reads) sequencing platform at the Auburn University’s Biological Sciences Department. Quantification of final library before loading on MiSeq sequencer was performed using the Kapa quantification kit (RT-PCR) for next generation sequencing with the Illumina platform (Kapa Biosystems, Wilmington, MA USA). Sequence reads were quality filtered and assembled using CLC Genomics Workbench 8.0.1 (CLCbio, Aarhus, Denmark) (K. U. Kim et al., 2013), SPAdes 3.6 (Bankevich et al., 2012) and Velvet 1.2 (Zerbino & Birney, 2008). QUAST β (http://quast.bioinf.spbau.ru/) (Gurevich, Saveliev, Vyahhi, & Tesler, 2013a, 2013b) was used to check for the quality of the assemblies and determine % G+C content of genomes. FASTA-formatted genomic sequences of closely related Staphylococcus genomes were obtained from the RefSeq database on GenBank and uploaded to PATRIC for annotation.

Protein coding genes in genomes were identified using GeneMarkS (Borodovsky & Lomsadze, 2014). Non-coding RNA prediction was achieved using RNAMMER 1.2 online server (Lagesen et al., 2007). Predictions for tRNA and tmRNA genes were done with the ARAG ORN tRNA and tmRNA prediction program (Laslett & Canback, 2004). Functional annotation of the protein gene models was achieved using multiple bioinformatic softwares including the RAST server (Overbeek et al., 2014), PATRIC (Wattam et al., 2014) and IMG/M (Markowitz et al., 2012). Annotation for antibiotic resistance genes was achieved using PATRIC via BLASTP sequence homology search from the Antibiotic Resistance Genes Database (ARDB) (Liu & Pop, 2009) and the Comprehensive Antibiotic Resistance Database (CARD) (McArthur et al., 2013) databases. To remove database-based redundancy, replicates were removed.

Analysis for occurrence of metal resistance genes (MRG) was performed using BLASTX against the Antibacterial Biocide and Metal Resistance Genes Database (MRDB) database with an e-value cutoff of 0.01 (Altschul et al., 1997). To achieve this, the BaCMet experimentally confirmed database of MRDB was used (Pal, Bengtsson-Palme, Rensing, Kristiansson, & Larsson, 2014).

Using the 16S rRNA gene sequences of strain AOAB and closely related *Staphylococcus* representatives were structurally aligned using SSU-ALIGN v0.1.1(Nawrocki, 2009) and used for reconstruction of neighbor-joining tree. Separate sequence alignments was done using ClustalW algorithm in MEGA7 (Kumar, Stecher, & Tamura, 2016) for unrooted neighbor-joining tree (Supplementary Figure S3).

FASTA sequences of housekeeping genes from the closest species were obtained from GenBank. Concatenated DNA sequences from six marker genes (16S rRNA, tuf, sodA, dnaJ, hsp60 and rpoB) were used for MLSA. Sequences were aligned using MUSCLE (Edgar, 2004). The evolutionary history of strain AOAB was inferred by using the Maximum Likelihood method based on the Jukes-Cantor model (Jukes & Cantor, 1969). Evolutionary analyses were conducted in MEGA7 (Kumar et al., 2016).

To infer genomic distance between strain AOAB and the closest species, pairwise average nucleotide identity (ANI) was computed using IMG/M system (Varghese et al., 2015). With a species cut off set at 96%, this method has been found to be robust in delineating bacteria based on genome sequence data (M. Kim, Oh, Park, & Chun, 2014).

*In-silico* DNA-DNA hybridization (DDH) was achieved using GBDP (Genome Blast Distance Phylogeny), which reliably infers genome-to-genome distances by utilizing Genome Blast Distance Phylogeny using logistic regression (Meier-Kolthoff, Auch, Klenk, & Goker, 2013).

## Results and Discussion

*Staphylococcus* sp. strain AOAB as determined by scanning electron microscopy (SEM) (Hitachi) depicted numerous clumps of coagulase-negative, vancomycin and oxacillin-susceptible bacterium (Figure 1). *Staphylococcus* sp. strain AOAB grown on LB and Mannitol agar at 30 °C produced yellow, medium (2-3mm), round, smooth colonies after 3 days (Supplementary Material 2) after t periods of growth. The isolate is catalase-positive and coagulase-negative. The polar lipid profile consisted of glycolipid, phospholipid, phosphatidylglycerol and diphosphatidylglycerol (Supplementary Figure S5), typical of *Staphylocccus* species (Nahae, Goodfellow, Minnikin, & Hajek, 1984). But unlike *S. warneri* and other *Staphylococcus* species (Nahae et al., 1984), *strain* AOAB did not have detectable β-gentiobiosyl diacylglycerol (Supplementary Figure S5) as part of its polar lipid profile.

**Figure 1:**
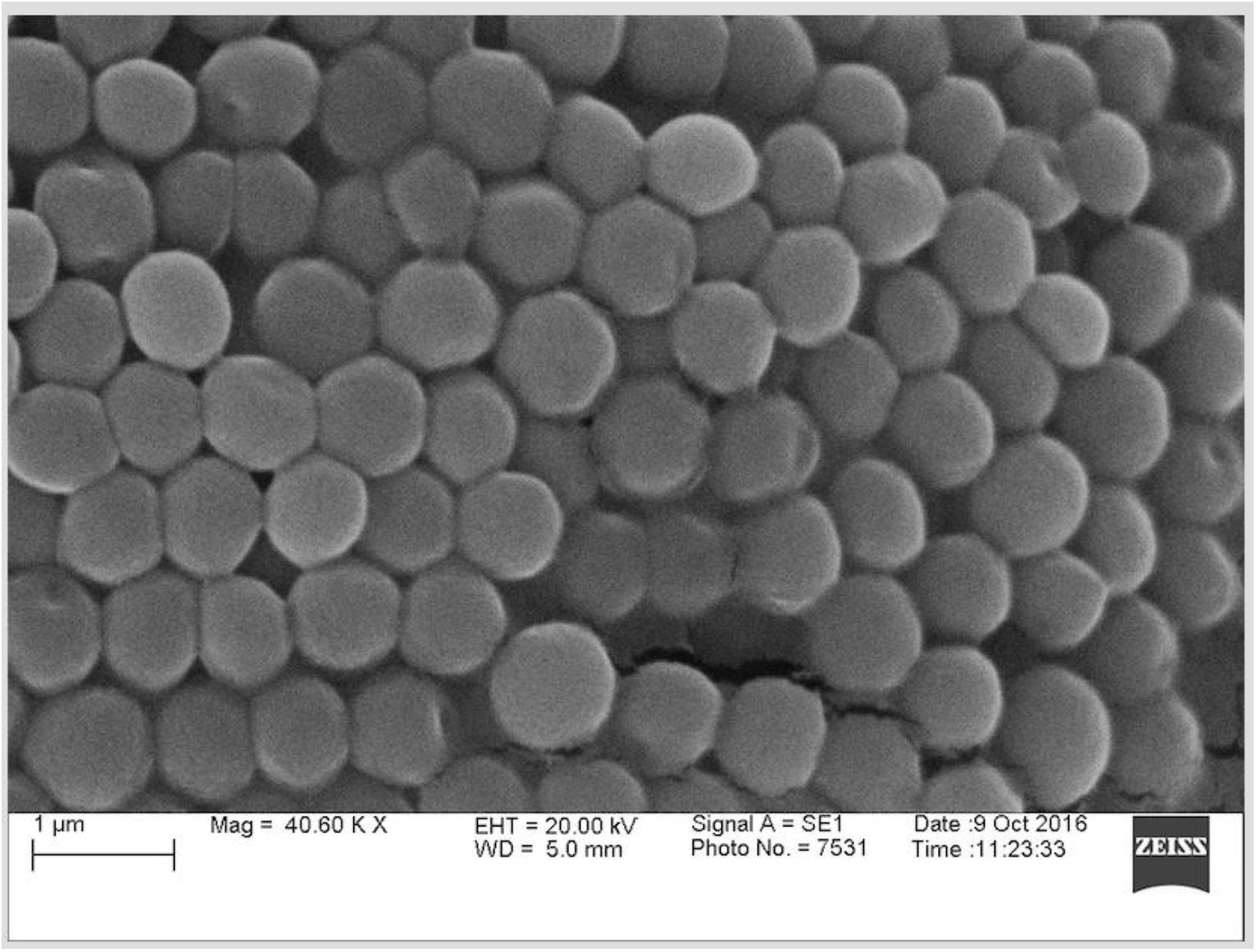
*Staphylococcus* sp. strain AOAB as determined by scanning electron microscopy (SEM) (Hitachi) depicted numerous clumps.

The presence of fatty acids, ai-C_15 : 0_, i-C_15 : 0_, i-C_17 : 0_, ai-C_17 : 0_, confirmed Genus of isolates as *Staphylococcus* (Supplementary Figure S4). Cluster analysis of the fatty composition of our isolate based on Euclidean distance revealed that our isolate does not belong to any known species (Figure 4) as MIDI dendrogram software places same species link at about 10 Euclidian Distance (http://www.midilabs.com/fatty-acid-analysis). The presence of menaquinone (MK-7) in the cytoplasmic membrane helped confirm the isolated as coagulase negative (Heß & Gallert, 2015).

Disk diffusion confirmed resistance of *S. mneniopsis* AOAB against the penicillins (Penicillin and Ampicillin), fluoroquinolones (Ciprofloxacin, Nalidixic acid), a polypeptide (Bacitracin) and an aminoglycoside (Kanamycin) (Table 2). Also, strain AOAB was revealed to be susceptible to vancomycin, oxacillin, tetracycline, Amoxicillin/clavulanic acid and chloramphenicol (Table 1). *S. pasteuri* strains have mixed results against tetracycline (Chesneau et al., 1993).

**Table 2:** Phenotypic characterization investigation using carbohydrate fermentation with the API CH50 systems (bioMe’rieux)

| Test/Characteristic | *S. mnemiopsis* | *S. pasteuri* | *S. warneri* | *S. hominis* | *S. lugdunensis* | *S. haemolyticus* |
| --- | --- | --- | --- | --- | --- | --- |
| GAL | ? | + | - | + | + | + |
| GLU | + | + | + | + | + | + |
| FRU | + | + | + | + | + | - |
| MNE | - | - | - | - | + | - |
| MAN | + | + | + | - | - | - |
| SOR | - | - | - | - | - | - |
| NAG | - | - | - | + | + | - |
| MAL | + | + | + | + | + | + |
| LAC | - | - | - | + | + | ? |
| SAC | + | + | + | + | + | + |
| TRE | + | + | + | + | + | + |
| DXYL | ++ | - | - | ND | ND | ND |

Only the tests with a positive result were included. GAL, D-galactose; GLU, D-glucose; FRU, D-fructose; MNE, D-mannose; MAN, D-mannitol; SOR, Sorbitol; NAG, N-acetylglucosamine; MAL, D-maltose; LAC, D-lactose SAC, Sucrose; TRE, D-trehalose; DXYL, D-Xylose

Phylogenetic analysis using the 16S rRNA gene using neighbor-joining method indicated that the isolate was closely related to *S. pasteuri* and *S warneri* (Figure 2). Further phylogenetic analysis using MLSA (Figure 3) clustered the isolate as novel bacteria. Both ANI and DDH using genome sequence data confirmed delineation of the isolate as a novel species. None of the closest species met the ANI cut off of 96% or the DDH cut off of 70% (Table 3).

**Table 3:** Pairwise (ANI) and DDH between *S. mnemiopsis* AOAB (ANI2) and closely related species. Entries represent per cent values.

| **Species** | **ANI1>2** | **ANI2>1** | **DDH** |
| --- | --- | --- | --- |
| *S. warneri* SG1 | 85.02 | 84.93 | 27.80 |
| *S. pasteurii* SP1 | 84.79 | 84.58 | 27.60 |
| *S. hominis* C80 | 79.62 | 79.59 | 22.40 |
| *S. haemolyticus* JCSC1435 | 79.52 | 79.31 | 22.60 |
| *S. saprophyticus* ATCC 15305 | 78.03 | 77.66 | 21.90 |
| *S. aureus* MRSA252 | 78.85 | 78.73 | 22.70 |
| *S. epidermidis* ATCC 12228 | 79.4 | 79.36 | 19.00 |
| *S. lugdunensis* HKU09-01 | 78.21 | 78.04 | 22.0 |
| *S. hominis* VCU122 | 79.62 | 79.59 | 22.40 |

**Figure 2:**
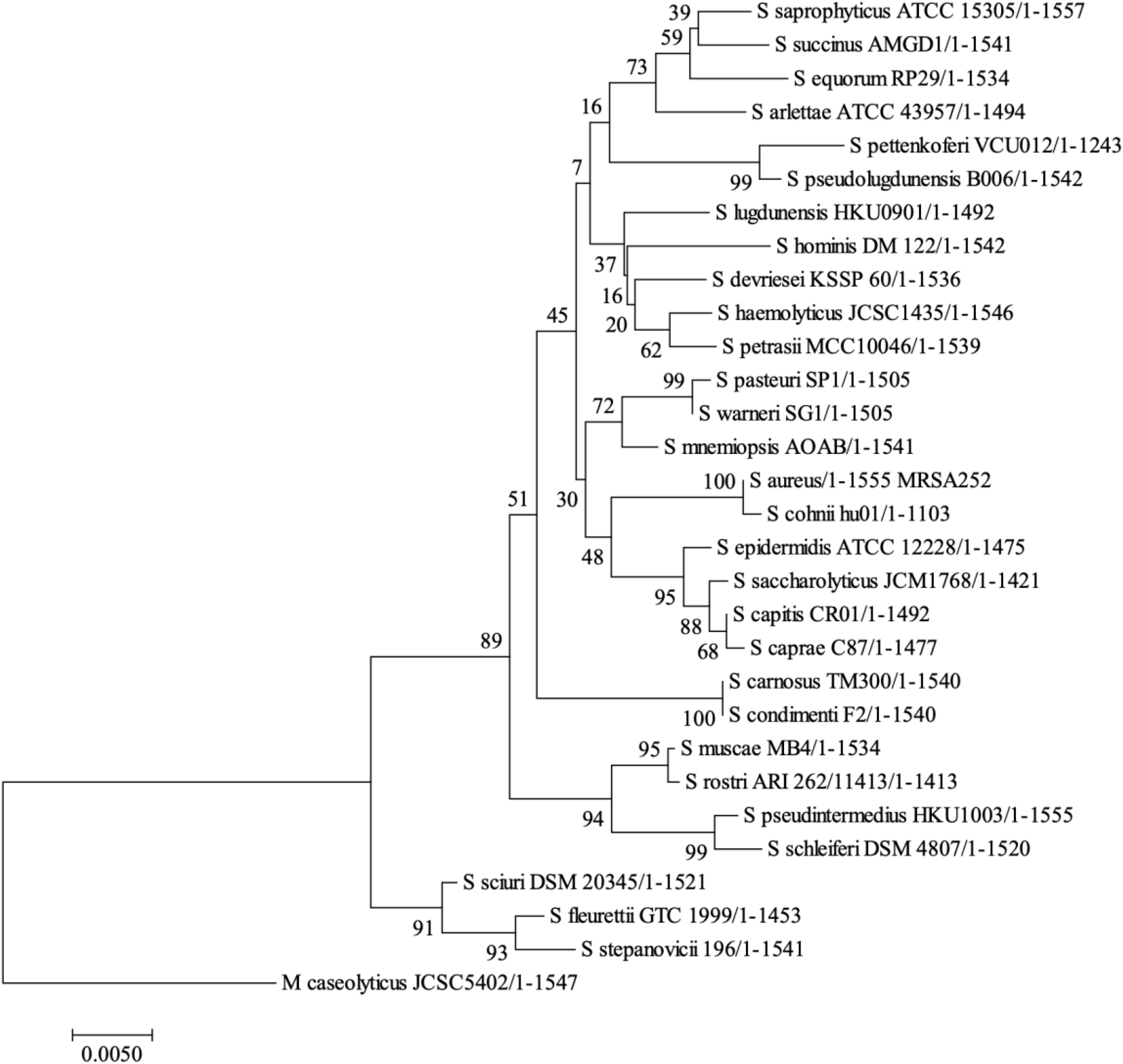
Neighbor-joining tree based on SSU-ALIGN for 16S rRNA gene sequences of strain AOAB with closely related *Staphylococcus* species. Numbers at nodes indicate the percentage of bootstrap support based on 1500 replications. Rooting was done using *Macrococcus caseolyticus* for discriminatory purposes.

**Figure 3:**
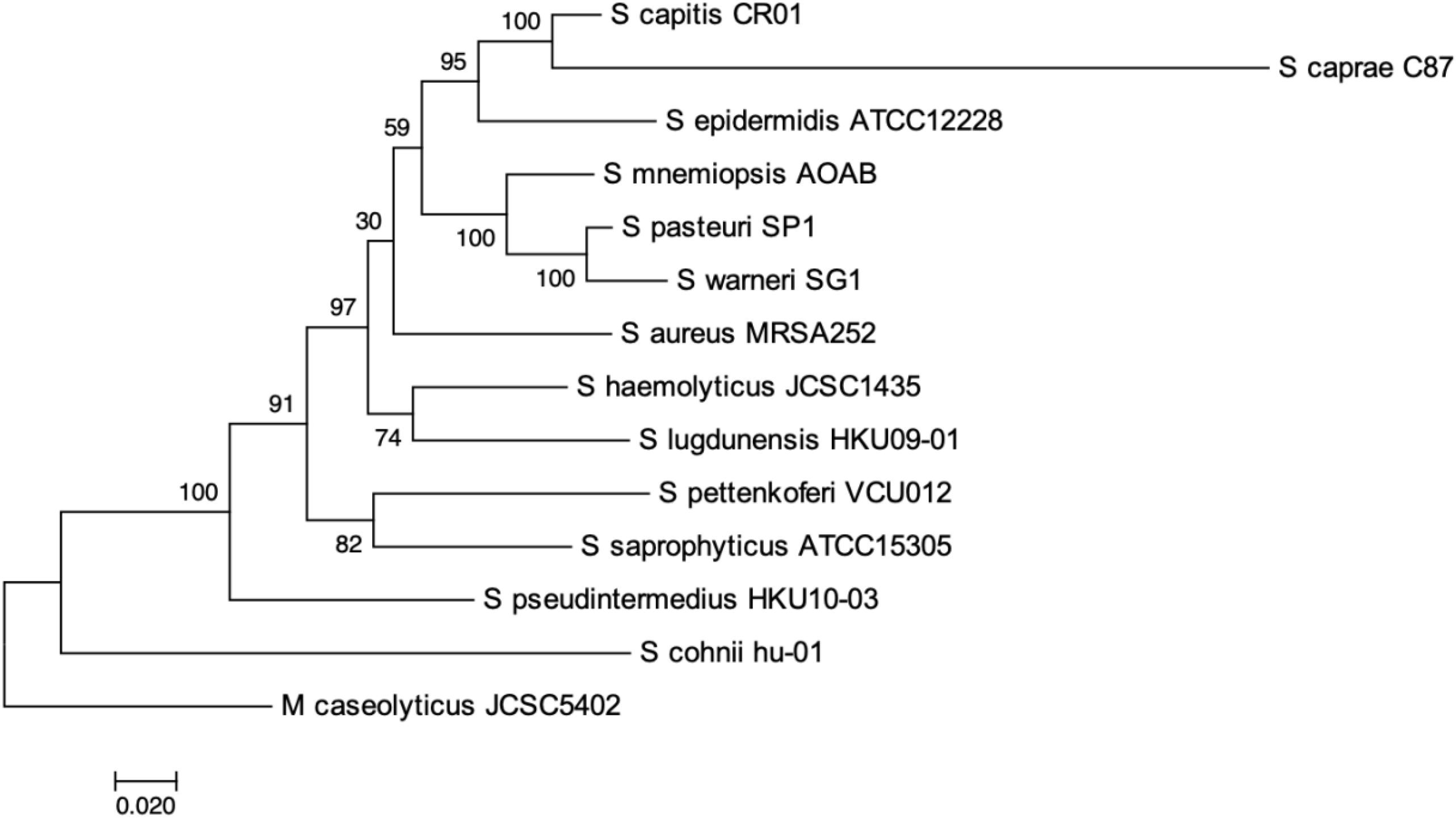
Evolutionary history of strain AOAB inferred by using Maximum Likelihood method based on Jukes-Cantor model. Evolutionary tree was constructed using MEGA7. The tree is drawn to scale. Numbers on the nodes indicate bootstrap values as a percentage based on 1000 replications with branch lengths measured in the number of substitutions per site. The tree was based on concatenated MLSA of 16S rRNA, *tuf, sodA, dnaJ, hsp60* and *rpoB* gene sequences.

**Figure 4:**
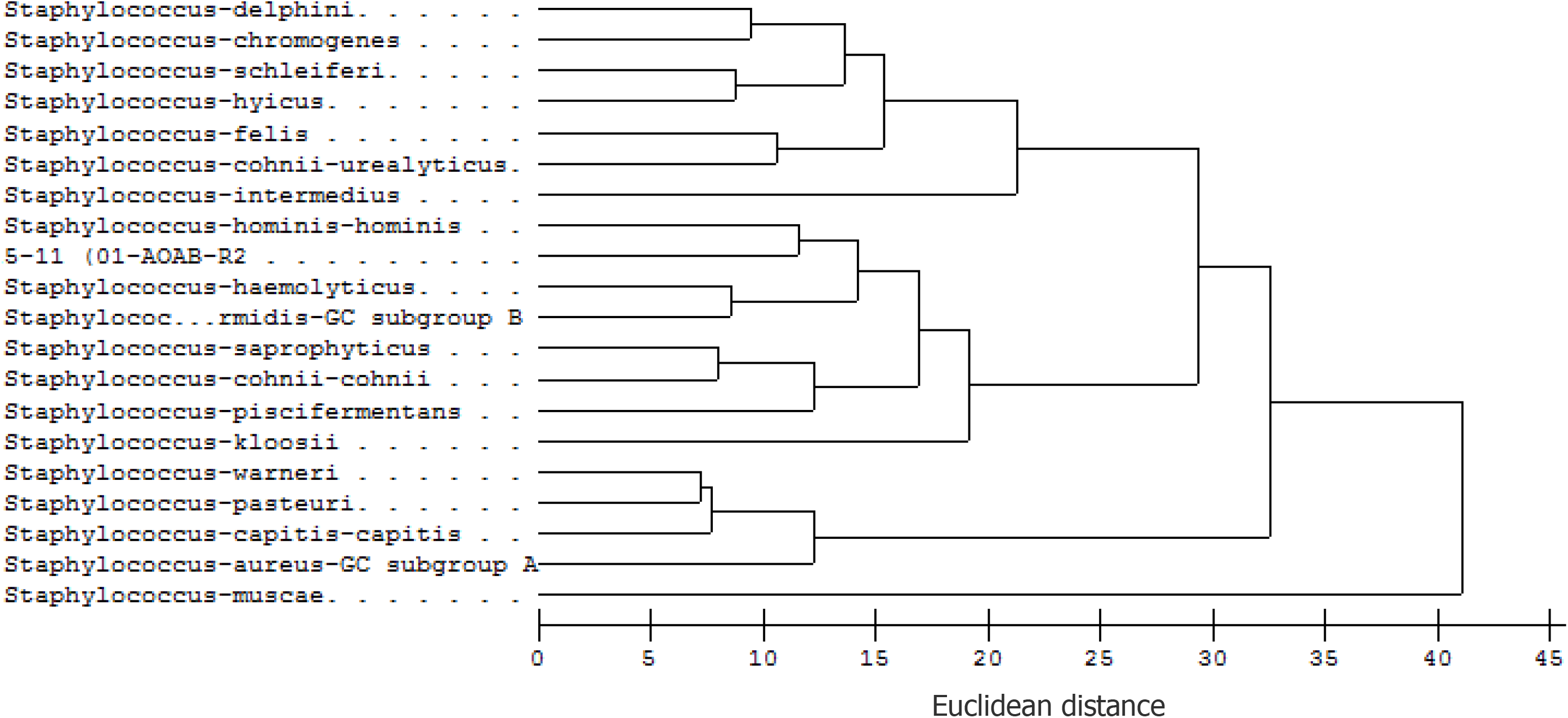
A dendrogram showing GC fatty acid profile similarities with 19 other *Staphylococcus* species.

Whole genome sequencing yielded a total of 905,410 paired reads for strain AOAB. The PATRIC annotated genome size was 2,617,061 bp. A number of genomic features differentiated it from the two closest relatives. Unlike *S. mnemiopsis, S. pasteuri* was experimentally confirmed to be susceptible to kanamycin (Chesneau et al., 1993), which helped in discrimination between the two species (Table 4).

**Table 4:** Genome characteristics of *S. mnemiopsis* AOAB and close relatives.

| Feature | *S. mnemiopsis*<br>AOAB | *S. warneri* SG1 | *S. pasteurii* SP1 |
| --- | --- | --- | --- |
| Size (bp) | 2,617,061 | 2,560,716 | 2,559,946 |
| % G+C | 32.13 | 32.73 | 32.71 |
| CDS | 2,695 | 2,435 | 2,457 |
| rRNA | 9 | 16 | 16 |
| tRNA | 59 | 60 | 48 |
| tmRNA | 1 | 1 | 1 |
| Hypothetical proteins | 595 | 539 | 559 |
| Proteins functional assignments | 2,100 | 1,896 | 1,898 |
| Proteins with EC number assignments | 770 | 739 | 719 |
| Proteins with GO assignments | 695 | 737 | 728 |
| Proteins with Pathway assignments | 630 | 579 | 559 |
| Proteins with FIGfam assignments | 2,294 | 2,214 | 2,194 |
| Kanamycin | resistant | ND | susceptible |
| Novobiocin | susceptible | susceptible | susceptible |

Genomic characterization of strain AOAB revealed that the isolate harbors antibiotic resistance determinants (virulence factors, antibiotic resistance and drug targets and heavy metal resistance genes) (Table 5). In addition to antibiotic resistance genes, MRG-like sequences were detected in the genome of strain AOAB. The most dominant ones were copper (22), arsenic (11) and zinc (4) metal resistance genes. *S*.*warneri* had copper (12), zinc (7) cadmium (7) and arsenic (5) as the dominant metal resistance genes and for *S. pasteuri* SP1, it was copper (12), zinc (7), cadmium (6) and arsenic (5), thus aiding in further discrimination. This study forms the first report of *Staphylococcus* isolation from the stomodeum of *M. leidyi*. The results also suggest that *M. leidyi*, a notoriously invasive zooplankton harbors culturable but previously uncharacterized *Staphylococcus* species, some of which harbor antibiotic resistance genes. Future efforts will involve investigation of host-microbe interaction.

**Table 5:**
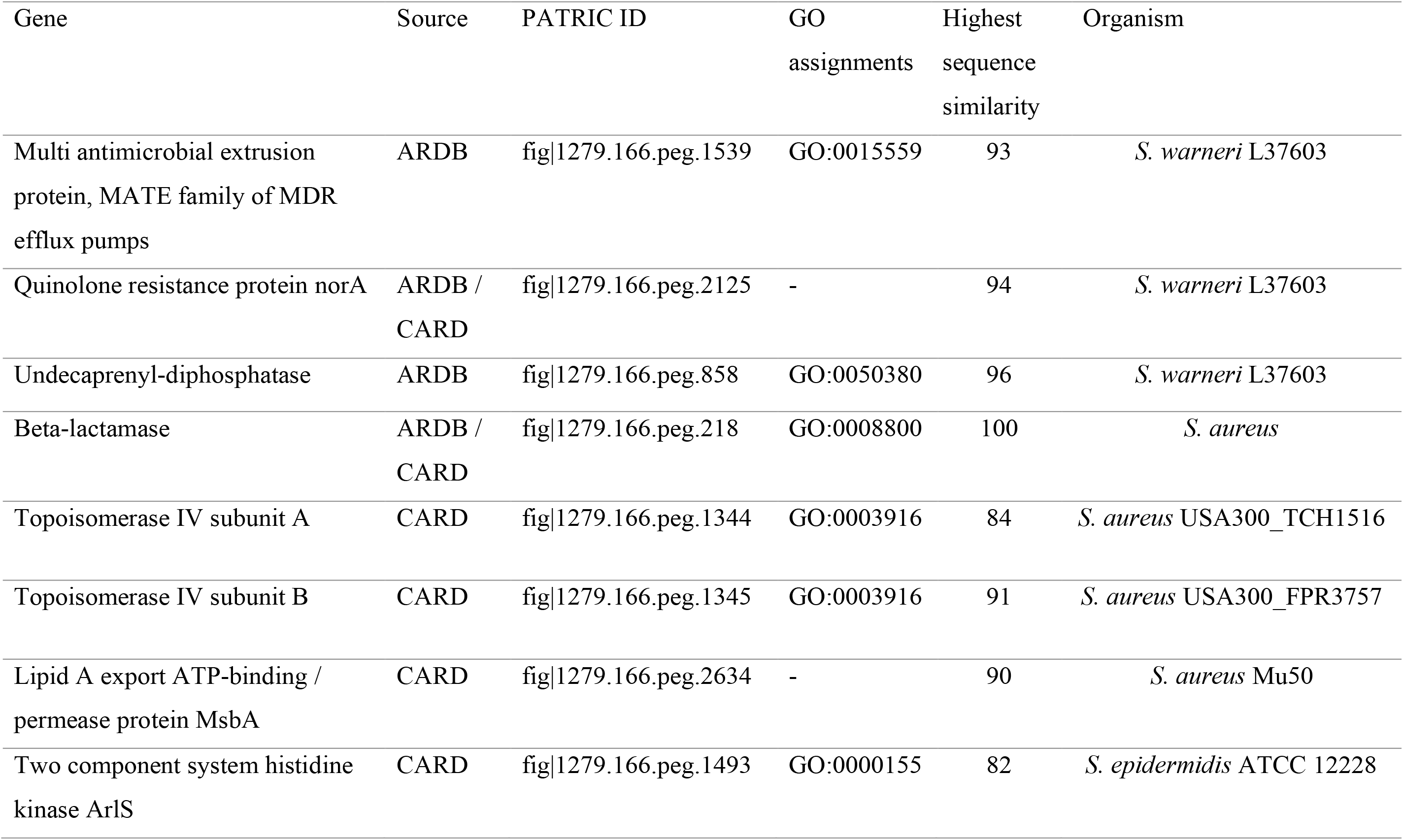
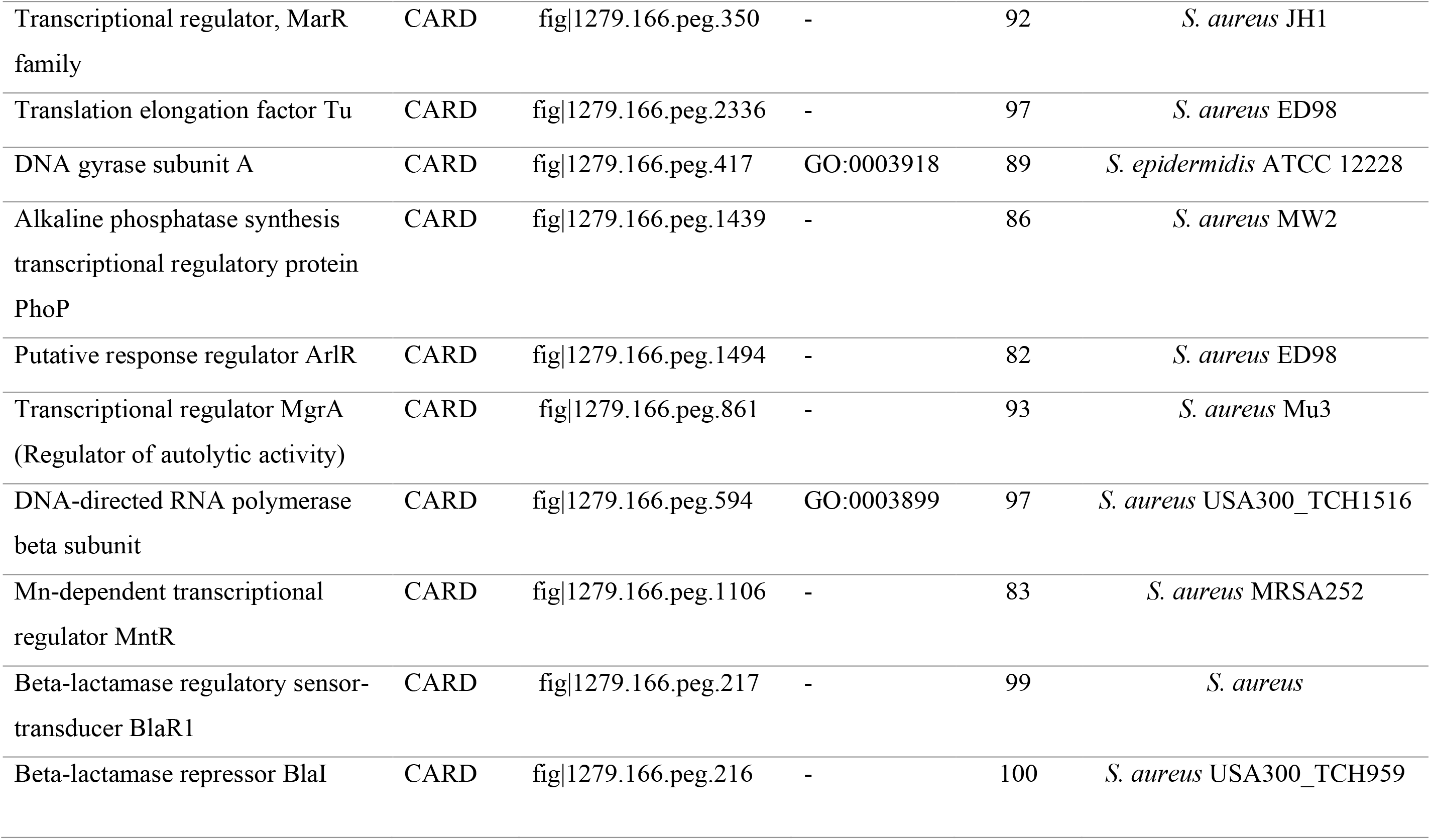
Antibiotic resistance genes from *S. mnemiopsis* AOAB (PATRIC Genome ID: 1279.166). Entries represent number of genes of the identified type.

## Supporting information

https://doi.org/10.6084/m9.figshare.13551431.v1

## Data Availability

The Whole Genome Shotgun project has been deposited at DDBJ/ENA/GenBank under the accession LZFL00000000. The version described in this paper is version LZFL01000000.TheSRA accession number is SRP076995, 16S rRNA gene (KU497670.1), tuf (LC158075), sodA (LC158854), rpoB (LC158855), hsp60 (LC158856) and dnaJ (LC158857). RAST server IDs are: Strain AOAB (6666666.127652), S. *pasteuri* SP1 (6666666.123700), *S. warneri* SG1 (6666666.123634). PATRIC Genome IDs: Strain AOAB (1279.166), *S. pasteuri* SP1 (1276282.3), *S. warneri* SG1 (1194526.3). Supplementary can be accessed via: https://doi.org/10.6084/m9.figshare.13551431.v1

## Acknowledgements

Technical and material assistance from We thank Dr. Lee Zhang, Dr. Scott Miller, Dr. Stephen Kempf (Auburn University) and Srijak Bhatnagar (UCDavis) for their technical and material assistance. We thank Prof. Aharon Oren (The Hebrew University of Jerusalem, Israel) for his technical advice.

## Funding

This work was supported in part by an NSF/EPSCoR PhD fellowship (NSF EPS-115886) and AU-Cell & Molecular Biosciences (CMB) Doctoral Graduate Research Fellowship to R.M.M. The study was also partly funded by USDA-Hatch 370225-310100 (AGM, MRL) and NSF0348327 (AGM).

## Competing interests

The authors have no competing interests to declare

## Abbreviations and key words

ANI: Average Nucleotide assembly
MLSA: Multilocus Sequence Analysis
DDH: DNA-DNA Hybridization

## Notes

https://doi.org/10.6084/m9.figshare.13551431.v1

### Competing Interest Statement

The authors have declared no competing interest.

