## Supplementary material for "Comparative genomic analysis and characterization of *Staphylococcus* sp. AOAB, isolated from a notoriously invasive *Mnemiopsis leidyi* gut revealed multiple antibiotic resistance determinants": https://doi.org/10.6084/m9.figshare.13551431.v1

**Supplementary Data**

Richard M. Mariita[1, 2], Mohammad J. Hossain [3], Anthony G. Moss[1]

[1]College of Science and Mathematics, Biological Sciences Department, Auburn University, AL 36849, USA

[2]Microbial BioSolutions, 33 Greene Street, Troy, NY, 12180 USA

[3]The Department of Biological Chemistry, The Johns Hopkins University School of Medicine, 725 N. Wolfe Street, Baltimore, MD 21205-2185, USA

Address correspondence to:

AGM:

RMM:

Supplementary Methods S1: Culture medium for enrichment, isolation of ctenophore endobacteria

Supplementary Figure S2: Growth of yellow pigmented *Staphylococcus mnemiopsis* AOAB on LB agar after 3 days at 30°C

Supplementary Figure S3: Unrooted neighbor-joining tree based on ClustalW alignments for 16S rRNA gene sequences of *S. mnemiopsis* AOAB with closely related *Staphylococcus* species. Numbers at nodes indicate the percentage of bootstrap support based on 1500 replications.

Supplementary Figure S4: Cellular fatty acid composition for *Staphylococcus mnemiopsis* AOAB using gas chromatography (chromatograph was fitted with a 5% phenyl-methyl silicone capillary column). Peak areas (a) and standard deviation (b) on percentage composition identified 17 fatty acids.

Supplementary Figure S5: Polar lipids analysis of *Staphylococcus mnemiopsis* AOAB (= DSMT102048 =NRRL B-65367T) using two-dimensional silica gel TLC (Macherey-Nagel Art. No. 818 135)

Supplementary Methods S1

The enrichment medium that was used was based on Anacker and Ordal with modifications as described by Figueiredo et al and Pilarski et al. (Anacker & Ordal, 1955; Figueiredo et al., 2005; Pilarski, Rossini, & Ceccarelli, 2008). Specifically, an infusion of ctenophore *Mnemiopsis* peptone was prepared using pieces of ctenophore (15 g of dry frozen weight) in 1L of distilled water. Ctenophore tissues were ground (pestle and mortar) and boiled for five minutes. The liquid was allowed to settle for thirty minutes at room temperature (25 ˚C) and filtered through 0.2 µm filters (Whatman® membrane filters nylon). The infusion was added to broth and solid mannitol growth media after autoclaving at 120°C for 15 minutes.

The enrichment broth was composed of select peptone (Life technologies, Cat. No. 30392-021) 3 g/L, yeast extract (Sigma, No. Y-4000), 5 g/L, sodium acetate (Fisher Scientific), 0.02 g/L 1, sodium chloride (Fisher Scientific), 0.02 g/L, ctenophore *M. leidyi* peptone, 100 mL/L and mannitol salt agar (BD and Sparks co., MD 21152 Ref. 211407-500g,) 27.75 g/L). Regular growth and isolation of *Staphylococcus* species was conducted using mannitol salt agar (BD and Sparks co., MD 21152 Ref. 211407-500g,) and LB agar (Sigma Aldrich).

Supplementary Figure S2

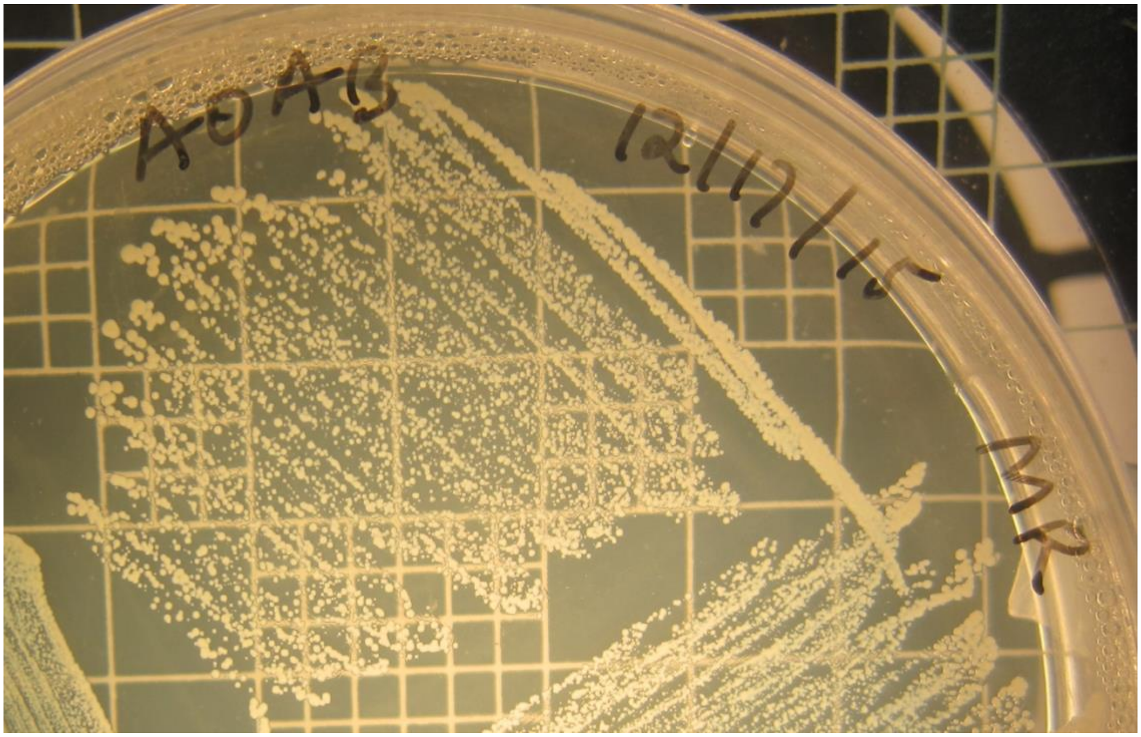

Supplementary Figure S3

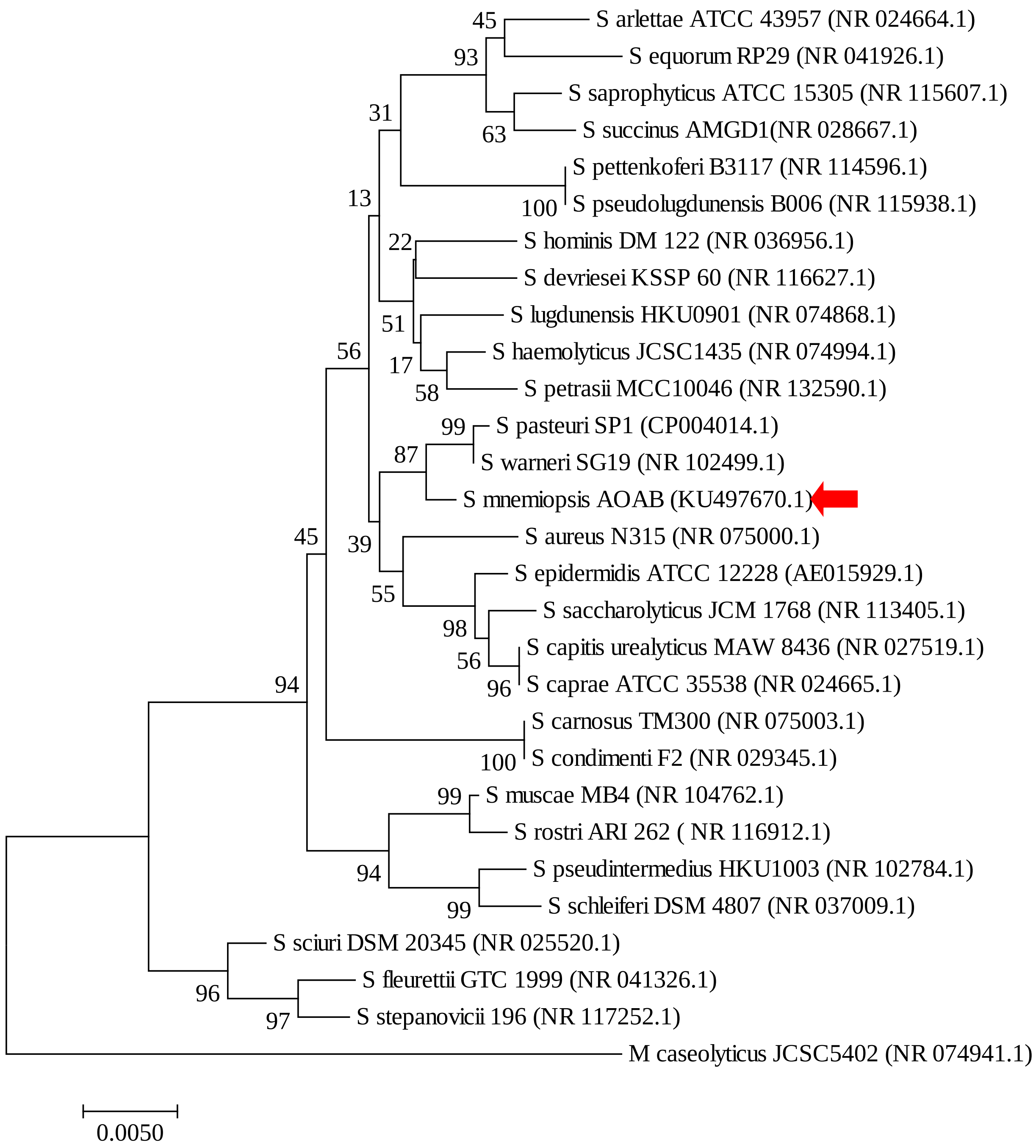

Supplementary Figure S4

a.

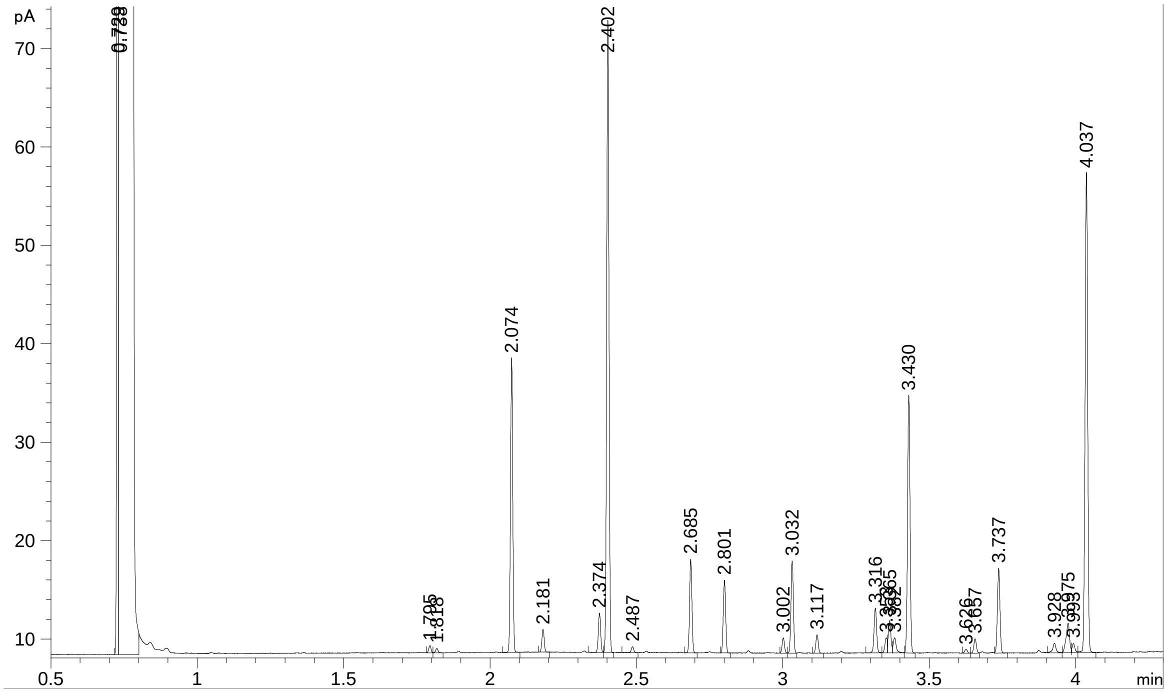


b.

| RT | Response | Ar/Ht | RFact | ECL | Peak Name | Percent |
|---|---|---|---|---|---|---|
| 0.7291 | 200492 | 0.005 | ---- | 6.6227 |  | ---- |
| 0.7376 | 1.191E+9 | 0.018 | ---- | 6.6813 | SOLVENT PEAK | ---- |
| 1.7949 | 543 | 0.008 | 1.027 | 12.6233 | 13:0 iso | 0.17 |
| 2.0741 | 5888 | 0.008 | 0.994 | 13.6277 | 14:0 iso | 1.83 |
| 2.1813 | 841 | 0.009 | 0.984 | 13.9997 | 14:0 | 0.26 |
| 2.3740 | 21254 | 0.009 | 0.969 | 14.6311 | 15:0 iso | 6.43 |
| 2.4030 | 134286 | 0.009 | 0.967 | 14.7261 | 15:0 anteiso | 40.52 |
| 2.6859 | 5319 | 0.009 | 0.949 | 15.6335 | 16:0 iso | 1.58 |
| 2.8010 | 10225 | 0.009 | 0.944 | 15.9991 | 16:0 | 3.01 |
| 3.0018 | 16624 | 0.009 | 0.936 | 16.6358 | 17:0 iso | 4.86 |
| 3.0329 | 44699 | 0.009 | 0.935 | 16.7343 | 17:0 anteiso | 13.04 |
| 3.1173 | 318 | 0.009 | 0.932 | 17.0019 | 17:0 | 0.09 |
| 3.3162 | 3574 | 0.009 | 0.927 | 17.6353 | 18:0 iso | 1.03 |
| 3.4304 | 39929 | 0.009 | 0.925 | 17.9987 | 18:0 | 11.53 |
| 3.6260 | 6156 | 0.009 | 0.923 | 18.6371 | 19:0 iso | 1.77 |
| 3.6571 | 10734 | 0.009 | 0.923 | 18.7384 | 19:0 anteiso | 3.09 |
| 3.7375 | 588 | 0.009 | 0.922 | 19.0010 | 19:0 | 0.17 |
| 3.9284 | 580 | 0.010 | 0.923 | 19.6368 | 20:0 iso | 0.17 |
| 4.0372 | 36243 | 0.010 | 0.923 | 19.9992 | 20:0 | 10.45 |
| 4.2534 | 525 | 0.009 | ---- | 20.7193 |  | ---- |

ECL Deviation: 0.001
Total Response: 337800
Percent Named: 100.00%

Reference ECL Shift
Total Named: 337800
Total Amount: 320321

Supplementary Figure S5

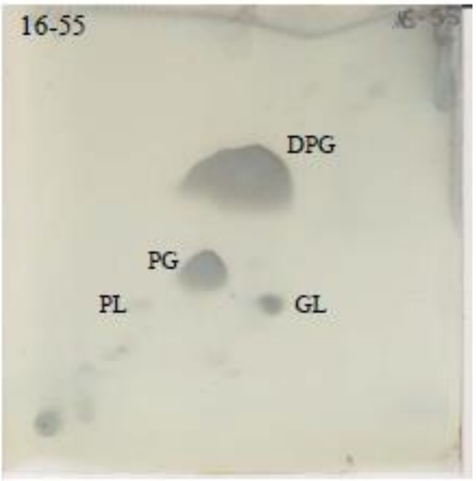


GL = Glycolipid

PL= Phospholipid

PG = Phosphatidylglycerol

DPG = Diphosphatidylglycerol
